# The evolutionary history of class I aminoacyl-tRNA synthetases indicates early statistical translation

**DOI:** 10.1101/2022.06.09.495570

**Authors:** Jagoda Jabłońska, Yao Chun-Chen, Liam M. Longo, Dan S. Tawfik, Ita Gruic-Sovulj

**Affiliations:** Department of Biomolecular Sciences, Weizmann Institute of Science, Rehovot, Israel; Earth-Life Science Institute, Tokyo Institute of Technology, Tokyo, Japan; Department of Chemistry, Faculty of Science, University of Zagreb, Zagreb, Croatia

**Keywords:** aminoacyl-tRNA synthetase, primordial translation, ancestral reconstruction, gene duplication

## Abstract

How protein translation evolved from a simple beginning to its complex and accurate contemporary state is unknown. Aminoacyl-tRNA synthetases (AARSs) define the genetic code by activating amino acids and loading them onto cognate tRNAs. As such, their evolutionary history can shed light on early translation. Using structure-based alignments of the conserved core of Class I AARSs, we reconstructed their phylogenetic tree and ancestral states. Unexpectedly, AARSs charging amino acids that are assumed to have emerged later – such as TrpRS and TyrRS or LysRS and CysRS – appear as the earliest splits in the tree; conversely, those AARSs charging abiotic, early-emerging amino acids, *e*.*g*. ValRS, seem to have diverged most recently. Furthermore, the inferred Class I ancestor (excluding TrpRS and TyrRS) lacks the residues that mediate selectivity in contemporary AARSs, and appears to be a generalist that could charge a wide range of amino acids. This ancestor subsequently diverged to two clades: “charged” (which gave rise to ArgRS, GluRS, and GlnRS) and “hydrophobics”, which includes CysRS and LysRS as its outgroups. The ancestors of both clades maintain a wide-accepting pocket that could readily diverge to the contemporary, specialized families. Overall, our findings suggest a “generalist-maintaining” model of class I AARS evolution, in which early statistical translation was kept active by a generalist AARS while the evolution of a specialized, accurate translation system took place.

**Significance:** Aminoacyl-tRNA synthetases (AARS) define the genetic code by linking amino acids with their cognate tRNAs. While contemporary AARSs leverage exquisite molecular recognition and proofreading to ensure translational fidelity, early translation was likely less stringent and operated on a different pool of amino acids. The co-emergence of translational fidelity and the amino acid alphabet, however, is poorly understood. By inferring the evolutionary history of Class I AARSs we found seemingly conflicting signals: Namely, the oldest AARSs apparently operate on the youngest amino acids. We also observed that the early ancestors had broad amino acid specificities, consistent with a model of statistical translation. Our data suggests that a generalist AARS was actively maintained until complete specialization, thereby resolving the age paradox.

## Introduction

Faithful translation of genetic information is at the core of extant life. Translation relies on highly accurate RNA-and protein-based catalysis, exemplified by ribosome and aminoacyl-tRNA synthetases (AARSs), respectively.^1,2,3,4^ How translation evolved from a simple beginning to the complex and accurate contemporary state has been the subject of multiple studies. ^5,6,7,8^ The consensus view is that translation developed gradually, by coevolution of multiple components.^7,9^ Early protein synthesis was based on so-called abiotic amino acids that were spontaneously formed^10^, and on some minimal, simple system and code that generated compositional rather than sequence-specific proteins (statistical translation)^11^. With time, additional biosynthesized amino acids were included, and AARSs, tRNAs, and the ribosome evolved further^12,13^ to allow accurate, gene-coded synthesis. However, beyond this general view, supporting evidence and details are in short supply. For example, attempts to reconstruct the primordial ribosome have indicated that the peptidyl transferase center represents an ancient RNA core.^14,15^ Yet, these studies did not reveal the functional characteristics of the primordial translation machinery.

The other major players, AARSs^16^ – which define the genetic code by activating amino acids and coupling them to the cognate tRNAs — have also been the subject of extensive evolutionary analyses^17–20^. AARSs are ancient enzymes whose amino acid specificity was generally established by the appearance of the last universal ancestor (LUCA)^12^. They are divided into two evolutionarily distinct classes (I and II)^21^ and each consists of the same 10 members in all domains of life (with lysyl-tRNA synthetase being the only exception to the class rule^22^). It has been proposed that the so-called synthetic domain of AARSs, where activation of the amino acid through the formation of an adenylate intermediate and its transfer to tRNA take place, emerged first^9^. Accordingly, attempts to reconstruct the evolutionary history of the synthetic domains^23,20^, as well as its primordial catalytic core^24,25^ have been made but not with the resolution needed to reveal the functional characteristics of the class I and class II^26^ early ancestral states.

To get deepen our understanding of AARSs evolutionary history, we have reconstructed the ancestral states of the synthetic domain with the goal of inferring evolution of AARS amino acid substrate selectivity over time. We opted for class I AARSs as the evolutionary and functional context in which Class I emerged is tractable. The synthetic domain of Class I AARSs belongs to the ancient HUP superfamily^27,28^, comprising predominantly ligases that use ATP to activate various substrates via a substrate-AMP intermediate. To provide a reliable structure-based sequence alignment we selected the most conserved catalytic core of the class I synthetic domain, which includes the amino acid and ATP binding sites. The inferred class I AARS tree shows unique features that promoted us to propose a new model of enzyme emergence. The existence of a well-defined evolutionary neighborhood allowed us to infer the properties of the early Class I ancestors, which appears to be a generalist, thereby supporting a statistical model of early translation.

## Results and Discussion

### Evolutionary background of Class I and Class II AARSs

We aimed to reconstruct the functional characteristics of primordial translation by inferring the properties of the early AARS ancestors. This goal demands a rigorous phylogenetic analysis based on a reliable alignment that, in the case of highly diverged sequences, must be guided by structure. Furthermore, reconstruction of evolutionary relationships at this resolution benefit greatly from the existence of a suitable outgroup for the structural and phylogenetic analysis. We found that Class I and Class II AARSs strongly differ in their evolutionary context. The origin of Class I AARS is well defined within the HUP’s phylogenetic tree, which indicates that PP-ATPases emerged first, and later gave rise to HIGH nucleotidyl pyrophosphate transferases (HIGH-NTPs, **Fig. 1**)^28^. HIGH-NTPs split into two subclades: Clade 1 comprises all Class I AARSs and Clade 2 includes the ATP/CTP-dependent ligases. The latter group show high structure and sequence similarity to Class I AARSs. Using Clade 2 as an outgroup, we succeed in the construction of the stable Class I AARS phylogenetic tree (**Fig. 1** and below), which is prerequisite for the ancestral reconstruction. In sharp contrast, for Class II AARSs we could not identify an evolutionary context within which these enzymes emerged (for details see **Supplementary Data**), as has been noted before^29^. Using the same approach (**Fig. S1**) as for Class I AARSs (detailed below), we could not achieve a stable Class II phylogeny (**Fig. S2**), presumably due to the lack of a suitable outgroup, making ancestor reconstruction intractable. Therefore, we focused our evolutionary analysis and reconstruction of the ancestral states only on Class I AARSs.

**Figure 1.**
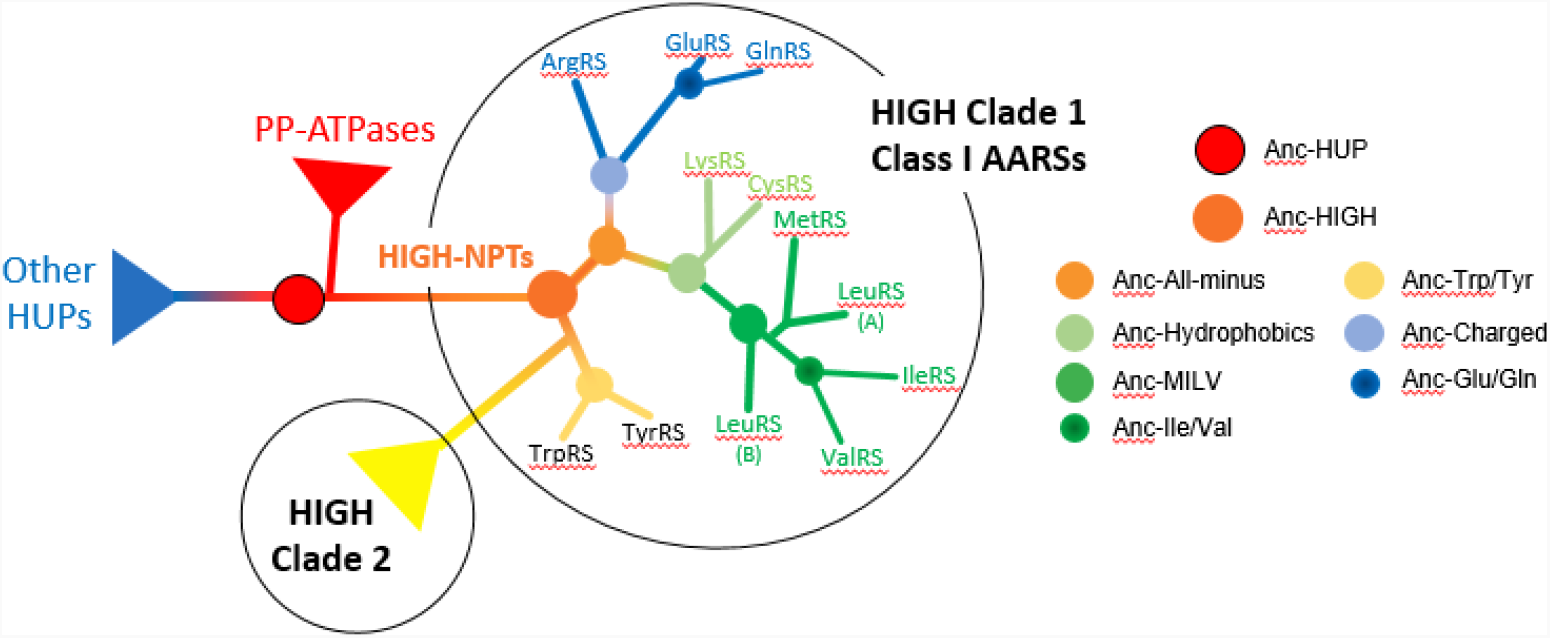
A schematic evolutionary tree of the HUP superfamily and Class I AARSs. Shown are the major splits inferred in our previous work^28^ and here, and the key ancestral nodes. The analysis described here focuses on HIGH nucleotidyl pyrophosphate transferases (HIGH-NTPs), which comprise two major Clades, 1 and 2, denoted by circles. Enzymes belonging to both of these clades possess a HIGH motif that mediates NTP binding and catalysis. This motif replaced the PP-loop motif observed in PP-ATPases, which represents the earliest HUP clade.

### Class I AARS tree revealed splits that coincide with the amino acid substrates physicochemical features

To provide a reliable structure-based sequence alignment for Class I AARS phylogenetic analysis, we extracted the most conserved structural segments of the HUP domain from a comparison of 24 AARS Class I structures (**Table S1**) and of seven Clade 2 enzymes that were used as outgroup (**Table S2**). HUPs adopt a specific fold characterized by five parallel β-strands and four α-helices, two at each side of the central β-sheet (**Fig. 2A**). The extracted core included all of these canonical elements (except one β-strand) and also the major functional elements. The core structures aligned well (**Fig. 2B; Datasets S1, Table S3**), as observed previously with a smaller set of structures^20^, allowing us to generate a manually refined, structure-guided sequence alignment (Fig. S3). As expected, this ‘seed’ alignment captured the two known Class I signature motifs (HIGH, KMSKS), as well as a GxD motif on the loop connecting β4 and α4. The latter mediates binding of the ribose moiety of ATP and is a hallmark of the HIGH-NPTs (present in both Clade 1, AARRs, and Clade 2)^28^.

**Figure 2.**
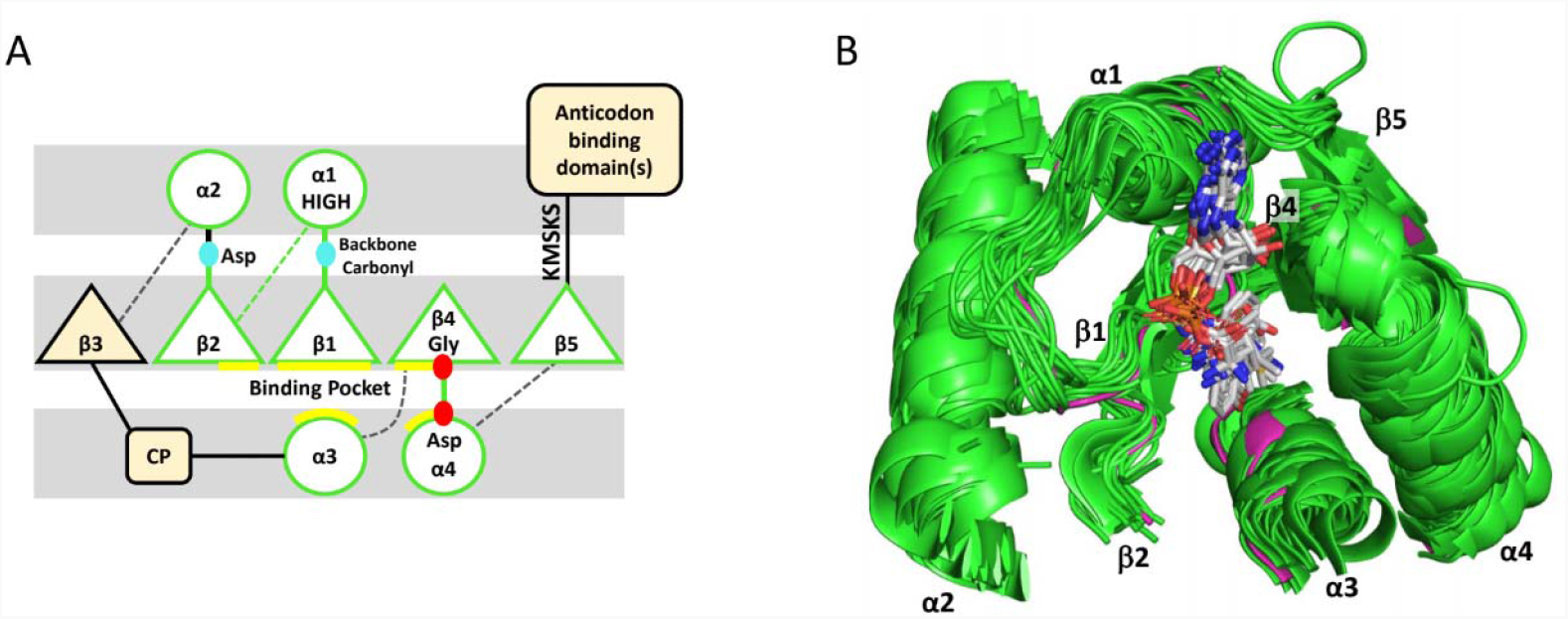
The aligned core of the Class I AARS synthetic domain. **A**. The HUP fold comprises parallel β-strands and α-helices connected via loops. The surface loops that make the active site are shown as full lines, while the ‘bottom’ connecting loops (dashed lines) have no known functional role. The surface loops vary in length and can include additional domains (the so-called CP fragments inserted between β3 and α3, and the anticodon binding domain that follows β5). The alignable core is marked by green lines and includes the key functional elements: The two known signature motifs, HIGH (top of α1) and KMSKS (the loop following β5); and the Asp residue within the β2 loop and another residue at the β1 loop that bind the α-amino group of the substrate amino acid (blue dots). The additional lines in yellow depict the amino acid substrate binding pocket, and the same color scheme is used throughout this paper. The red dots designate the ribose binding elements (the GxD motif, see text). **B**. An overlay of the core structures. Alignment of the extracted core segments of the selected Class I AARSs structures (N=24, **Table S1**) and the related HIGH-NTPs from Clade 2 (N=7, **Table S2**). The best overlap is of the β1-α1-β2 segment, while α-helices α3 and α4 show the highest variability. MetRS (PDB code 1pg0, colored magenta) was used as a template for defining the Class I core structure and for structural alignment of the Class I AARS cores (Table S3).

The AARS sequences from a set of 208 archaeal and 530 bacterial genomes, chosen to represent the prokaryote tree of life^30^ (**Datasets S2**), were added to the seed alignment. Phylogenetic trees were inferred from this alignment (**Fig. S4, Dataset S3**). In parallel, and similar to previous studies^23^, we performed an automated alignment of the entire HUP domain excluding the anticodon binding domains (**Dataset S4**) and use it to infer phylogenetic trees (**Dataset S5**). The anticodon binding domains are thought to have emerged later^9^, in agreement with our analysis, which indicated that the anticodon domains of Class I AARSs relate to at least three separate evolutionary emergences (**Table S1**).

The different trees, obtained with different methods (**Fig. S4, Dataset S3, S5**), converged to a consistent picture summarized in **Fig. 1**. Specifically, TyrRS and TrpRS appear to have diverged directly from the HIGH-NTPs ancestor (Anc-HIGH) alongside Clade 2. The next node denotes an ancestor from which all the remaining Class I enzymes diverged, Anc-All-minus, which in turn gave rise to two clades: the “Charged” clade, to which ArgRS, GluRS, and GlnRS belong, and the “Hydrophobics” clade. The latter includes the Met/Ile/Leu/ValRSs clade (the only unambiguously Class I clade assigned so far)^20,31,32^, CysRS (often assigned to this group)^31,32^, and somewhat unexpectedly also LysRS. This tree partially overlaps with the current divisions of Class I to subgroups (*e.g*., Met/Ile/Leu/ValRSs group together, and Tyr/TrpRSs)^20,31,32^ yet differs in other parts, foremost in Arg/Glu/GlnRSs being grouped together as well as Cys/LysRSs (which is also grouped with Met/Ile/Leu/ValRSs). Notably, the phylogenetic clades observed here coincide with the physicochemical characters of the amino acids: aromatics, charged, and hydrophobics, with the latter including the relatively hydrophobic Cys and the aliphatic Lys. But how did a single ancestor give rise to a repertoire of selective binding pockets covering range of physicochemical properties? To address this question, we examined the ancestral binding pockets.

### Class I early ancestors have wide and unspecialized amino acid substrate binding pockets

We inferred the ancestor sequences of the core segments (**Fig. S3, Fig. S5, Dataset S6, S7**) of each major AARS clade depicted in **Fig. 1** as well as each specialized AARS family. We then examined the ancestral states of the residues that comprise the amino acid substrate binding pocket in the contemporary enzymes (**Fig. 3, Table 1, Table S4)**. Not only is the amino acid binding pocket at the same location in all Class I enzymes, but largely the same positions mediate amino acid binding in all of the enzymes that diverged from Anc-All-minus. Two positions have a common role: Asp at the tip of β2 (β2B), which anchors the α-amino group, and a His or Gln at α4 (α4A), which generally anchors the carbonyl group of the carboxylate (**Fig. 3A**). The ancestral states of the common anchoring residues, β2B and α4A, are largely maintained from early nodes to the ancestors of specialized AARS families (**Table 1**). Other positions dictate the side-chain selectivity, and these vary between enzymes. In Anc-All-minus, and in its closest descendants Anc-Charged and Anc-Hydrophobics, the ancestral states of the positions that dictate the side chain selectivity indicate a wide pocket, with neither polar or charged interactions (as in the ancestors of ArgRS or Glu/GlnRS) nor a deep and hydrophobic pocket (as in the ancestors of the individual Met/Ile/Leu/ValRSs). Take for example β1A: in the Glu/GlnRS ancestor this residue is Arg, while in ArgRS ancestor it is Glu, thus contributing to selectivity toward either an anionic or a cationic amino acid substrate. In the early ancestors, this position is Ala, thus yielding a wider pocket devoid of charge preference. In general, all charged residues in the ancestors of contemporary enzymes are replaced with smaller uncharged residues (β2A, α3B, or α4C). Similarly, β1C, which is large and hydrophobic in ancestors of individual Met/LeuRS, or Pro in Ile/ValRS, is Gly in the early ancestors.

**Figure 3.**
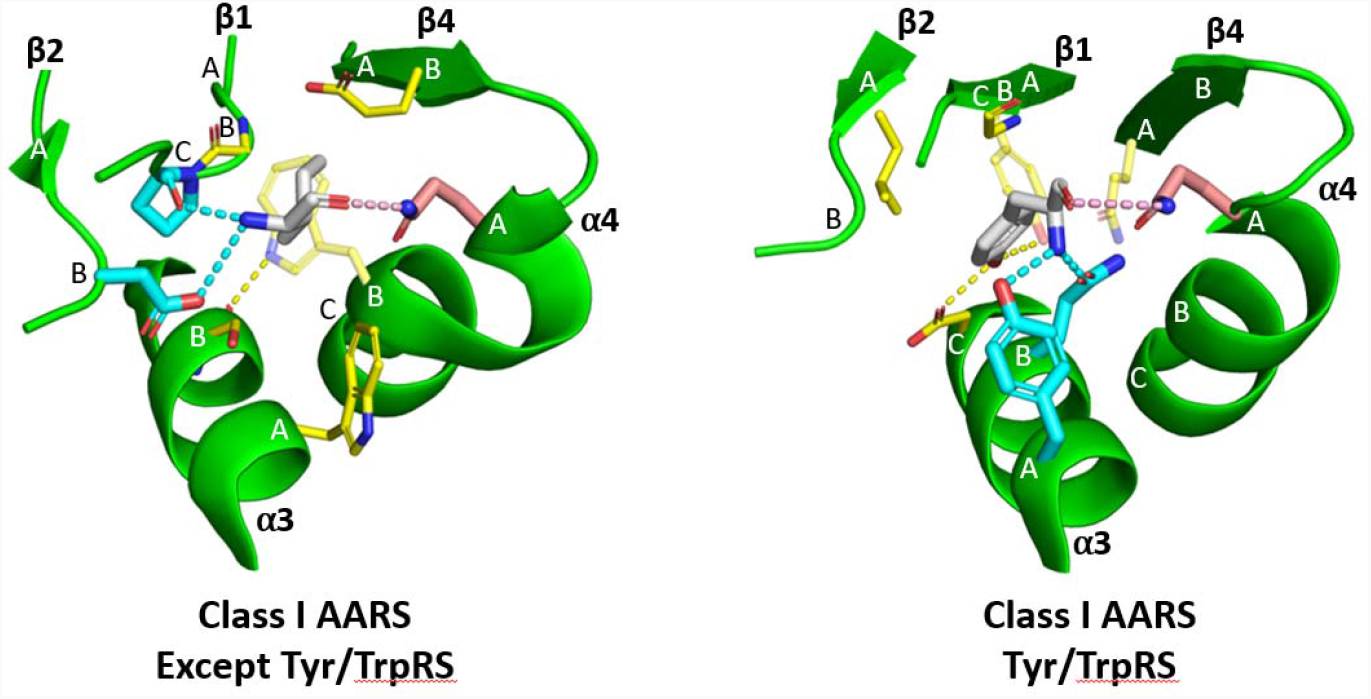
The amino acid substrate binding pockets of class I AARSs. The amino acid binding pocket of Class I AARSs resides between the β-sheet (β1, β2, and β4) and α3 and α4 (**Figure 2**). The residues that comprise the pocket wall are annotated by the secondary structure element on which they reside, and by their order from the N-to the C-terminus. The interacting side chains are shown only for the presented structures using the same color scheme as in **Table 1**: Residues interacting the substrate’s side chain are shown in yellow; residues interacting with the α-amino group are shown in blue; and residues interacting with the α-carboxyl group are shown in pink. The pocket’s composition, and the anchoring of the of the α-amino group, differs between Tyr/TrpRSs and all other class I AARSs. The latter are shown in **Panel A** (represented by IleRS, 1jzq) and the former in **Panel B** (represented by TyrRS, 1vbm).

**Table 1.** The amino acid substrate-binding pockets of Class I AARS family ancestors and selected earlier ancestral nodes. The ancestral states^i^ of the residues that comprise the amino acid substrate-binding pocket are bold and shaded yellow (side chain recognition), blue (α-amino group recognition), or orange (α-carboxyl group recognition). Double shading indicates interactions with multiple parts of the substrate. No shading indicates lack of interaction. Ancestral states of the early ancestors (prior specialization) are rendered without shading. Early ancestral states with smaller and neutral amino acids relative to the associated family ancestors are colored red. Charged early ancestral states are colored blue. The binding pockets were assigned from an analysis of contemporary AARSs.

|  | AARS family ancestors |  |  |  |  |  |  |  | AARS early ancestors |  |  |  | AARS family ancestors |  | AARS early ancestor |
| --- | --- | --- | --- | --- | --- | --- | --- | --- | --- | --- | --- | --- | --- | --- | --- |
|  | MetRS<br>(N672)<sup>ii</sup> | LeuRS<br>(N735<sup>iii</sup><br>N724) | ValRS<br>(N577<br>N571) | IleRS<br>N603 | CysRS<br>N787 | LysRS<br>Class I<br>(N769)<sup>iv</sup> | Glu/<br>GlnRS<br>(N927) | ArgRS<br>(N837) | Ile/Val<sup>v</sup><br>(N570) | "All-<br>hydropho<br>bic"<br>(N567) | "Charged<br>" (N836) | "All<br>minus"<br>(N566) | TrpRS<br>(N1077<br>N1121) | TyrRS<br>(N1040) | Trp/Tyr<sup>vi</sup><br>(N1039) |
| $\beta 1 A^{vii}$ | Thr | Leu<br>Thr | Val<br>Asp | Leu | Tyr | Ala | Arg | Glu | Thr | Ala | Ala | Ala | Leu<br>Met | Met | Val |
| $\beta 1 B$ | Pro | Met<br>Pro | Pro | Gly | Cys<br>Zn shell | – | – | Val | Pro | Val | Val | Val | – | – | |
| $\beta 1 C^{viii}$ | Ile | Phe | Pro | Pro | Gly | Gly | Ala | Ser | Pro | Gly | Gly | Gly | Gly | Gly | Gly |
| $\beta 1 D$ | Tyr | Tyr | Asn<br>Tyr | Tyr | Thr | Ser | Asn | Asn | Tyr | Asn | Asn | Asn | Arg<br>Met | Met | Met |
| $\beta 2 A$ | Gly | Gly | Gly | Gly | Asn | Val | Arg | Tyr | Gly | Ser | Ser | Ser | Ile<br>Val | Leu | Leu |
| $\beta 2 B$ | Asp | Asp<br>His | Asp | Asp | Thr | Asp | Asp | Asn | Asp | Asp | Asp | Asp | Ala<br>Leu | Ala | Ala |
| $\alpha 3 A$ | Trp | Phe<br>Leu | Trp<br>Val | Trp | Trp | Trp | Tyr | Tyr | Trp | Trp | Tyr | Tyr | Tyr | Tyr | Tyr |
| $\alpha 3 B$ | Ala | Ser | Ser<br>Lys | Ser | Cys<br>Zn shell | Trp | Ser | Asp | Ser | Ser | Ser | Ser | Gln | Gln | Gln |
| $\alpha 3 C$ | Asn | Tyr | Trp<br>Tyr | Ser | Met | Arg | Asp | His | Tyr | Asn | Asn | Asn | Asp | Asp | Asp |
| $\alpha 3$ D | Tyr | Phe<br>Ala | Pro<br>Val | Pro | Ser | Trp | Asp | Phe | Leu | His | His | His | Ile | Ile | Ile |
| $\beta 4$ A | His | Tyr<br>Arg | Leu<br>Arg | Tyr | Thr | Glu | Val | Asn | His | His | His | His | Val | Val | Val |
| $\beta 4$ B | Ile | Gly<br>Ser | Thr<br>Gln | Glu | Gly | Gly | Arg | Leu | Gln | Gly | Gly | Gly | Val | Val | Val |
| $\alpha 4$ A | Ile | His<br>Leu | Ile | Gln | His | His | His | His | Ile | His | His | His | Gln | Gln | Gln |
| $\alpha 4$ B | His | His | Trp | Trp | His<br>Zn shell | Ser | Thr | Tyr | Trp | His | His | His | His | Thr | His |
| $\alpha 4$ C | Trp | Ser<br>Phe | Met<br>Thr | Ser | Glu<br>Zn shell | Gly | Gln | Leu | Thr | Ser | Gln | Ser | Thr | Ala | Thr |

One possible caveat is that the tree topology whereby TyrRS and TrpRSs are the outgroup of Anc-All-minus (**Fig. 1**) could dictate ancestral states that would artificially make Anc-All-minus resemble Tyr/TrpRSs. However, the wide, non-discriminating pocket is a distinct characteristic of Anc-All-minus, and Anc-Trp/Tyr is inferred to have a distinctly different pocket that is very similar to the contemporary Tyr/TrpRSs. Furthermore, in Tyr/TrpRSs, the amino acid substrate is positioned differently compared to all other Class I enzymes. The α-amino group is anchored to α3 instead of β1 and β2, as in all enzymes that descend from Anc-All-minus; the residues that bind the amino acid side chain also differ (**Table 1; Fig. 3B**).

The different mode of substrate binding is in agreement with an independent emergence of Tyr/TrpRSs (**Fig. 1** and^33^).

### Unusual features of the class I AARS tree unveil a novel model of emergence of new enzymes

Independent emergence of Tyr/TrpRSs is also in agreement with the notion that these two aromatic amino acids were the latest addition to the canonical set of proteinogenic amino acids^34^. Late emergence is, however, in disagreement with Tyr/TrpRSs appearing as the earliest split in the Class I tree (**Fig. 1**). In principle, the early splits in phylogenetic trees correspond to sequences, or species, which are the closest contemporary representatives of the early ancestor(s). For example, the LUCA is best represented by contemporary archaeal and bacterial species that appear as the earliest splits in the tree of life, compared to humans that appear as one the latest splits. An early split may also result from inclusion of sequences that are highly diverged compared to all others (the so-called outgroup effect). However, this trend of early splits in the AARSs tree relating to later emerging amino acids also appears in inner parts of the tree: Lys/CysRSs are the earliest split from Anc-Hydrophobics, and MetRS appears as an earlier split than ValRS – despite Val being the simplest, most abiotically available amino acid among the ones used by Class I ^10^. Thus, by the traditional interpretation of trees, the Class I tree is inconsistent with the assumption that AARSs emerged by the order by which amino acids appeared – starting from the simplest abiotic one (Val) to the most complex ones (Tyr, Trp)^10,34^.

Further, the inferred Anc-All-minus pocket bears no hallmarks of a Val activating enzyme, thus ruling out the option that Class I enzymes diverged from a primordial ValRS. In fact, our inference indicates that Anc-All-minus possessed only one distinct selectivity feature - an Asp at the tip of β2 that binds the substrate’s α-amino group (**Table 1**). Jointly with the anchoring of the α-carbonyl to a His side chain on α4, Anc-All-minus seems to have evolved to bind and activate α-amino acids with L stereo-configuration. An entire range of β-amino acids, as well as α-hydroxy acids, with both L and D isomers, are seen in spark discharge experiments and extraterrestrial samples ^35,36^. Thus, Anc-All-minus seems to mark a key filtering step toward the regio-and stereo-selectivity of protein synthesis. However, with respect to the side chain, Anc-All-minus seems like a generalist with no particular selectivity. This generalist diverged by a series of duplications to ultimately give nine enzymes each specializing in one amino acid substrate. However, as noted above, the order of duplications (splits) is the opposite of what is expected from the order by which these amino acid substrates were recruited.

Broadly speaking, there are two different models describing how duplication drives the emergence of new proteins. The first follows Ohno’s model of *neo-functionalization*^37^. It assumes an ancestor with a given activity and selectivity (a specialist). Upon duplication, one copy maintains the original function while the other evolves towards a new one (**Fig. 4**). This mode of divergence did take place in AARSs, most evidently in the divergence of GlnRS from GluRS^17^ and also, by our analysis, in the divergence of IleRS from ValRS. Indeed, Anc-Ile/Val shows high resemblance to contemporary ValRSs, which suggests that activation of Ile was newly acquired after duplication (**Table 1, Fig. S4**). The Ile/Val ancestor likely activated Ile as a latent promiscuous activity which was then used as starting point for the divergence of IleRS. This divergence scenario is also in agreement with the order of appearance of the amino acids – Ile emerging after Val, and Gln after Glu^34^.

**Figure 4.**
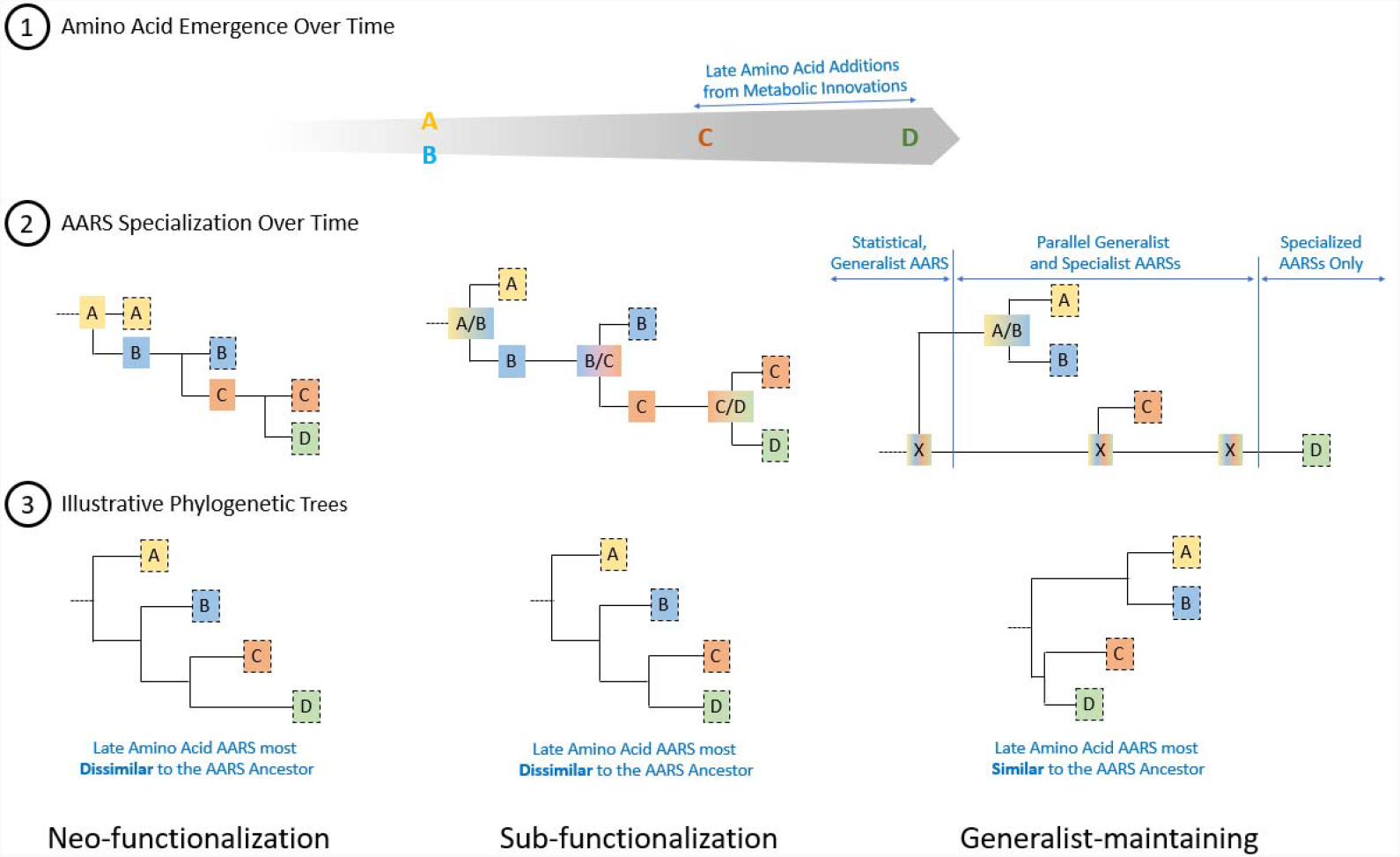
The alternative models for the emergence of new enzymes by duplication and divergence. Shown are schematic trees of divergence along evolutionary time (rather than sequence divergence) where new substrates, C and D, appear with time. By the sub-functionalization model, divergence is symmetric: the bifunctional ancestor is lost and both the original and the duplicated copy adopt new substrate selectivity (depicted by bifurcation). In contrast, in neo-functionalization, divergence is asymmetric - the original gene maintains the original function (substrate A) while the duplicate evolves a new function. This feature also underlies the new model of generalist-maintaining, except that the ancestor is a generalist that uses a wide range of substrates including A and B (denoted as X). Further, by this model (generalist-maintaining), the generalist ancestor is maintained while its duplicates give rise to specialized enzymes, and is lost only when the system becomes entirely specialized.

The alternative *sub-functionalization* model assumes a generalist, bi-or multi-functional ancestor. Upon duplication, the ancestral functions are split between the descendant duplicates, each specializing in one of the ancestral substrates (*e*.*g*., substrate A and B in the first duplication; **Fig. 4)**.^38^ Upon appearance of a new substrate whose intake provides an advantage (substrate C), one of the specialists would become bifunctional, and its duplication would allow the divergence of a new specialist. The divergence of Tyr/TrpRSs seems to follow this scenario. Their common ancestor looks nearly identical to the individual family ancestors and the contemporary enzymes. In particular, an Asp at the bottom of the binding pocket (α3 C) that dictates selectivity for the hydroxyl group Tyr on the one hand, and for the imine group of Trp on the other, is also present in Anc-Trp/Tyr (**Table 1, Fig. 2B**). Thus, Anc-Trp/Tyr likely emerged as a bifunctional enzyme incorporating both Tyr and Trp, and upon duplication diverged into two specialized enzymes.

Anc-all-minus, and the subsequent nodes (Anc-hydrophobics, Anc-charged) show clear hallmarks of a generalist, thus suggesting *sub-functionalization*. However, the tree topology suggests that only one copy led to specialization (like in *neo-functionalization*) while the other maintained the original generalist function. This observation provides the basis for a novel model of emergence dubbed generalist-maintaining (GM). By this model, the generalist enzyme forms were maintained and would subsequently serve as the starting point for AARSs to incorporate amino acids that appeared at later stages (**Fig. 4**), such as Lys/CysRSs, which diverged from Anc-hydrophobics (**Fig. 1**). That the ancestral active sites include late-emerging amino acids such as His (most notably in the HIGH motif) as well as Tyr and Trp (**Table 1**) suggests that the generalist enzyme(s) were still relevant close to LUCA, and possibly after.

### Concluding remarks

It has been suggested that translation evolved gradually from a simple, non-templated and less specific process to the coded and accurate systems of contemporary biology.^6,7,11,39^ These early beginnings likely included proto-tRNAs, a primitive ribosome built from the PTE core^14,40^, and a minimal AARS catalytic domain capable of activating amino acids and transferring them onto proto-tRNA.^6^ The latter was assumed to comprise the XCCA tail which later evolved to minihelices resembling the acceptor stem of contemporary tRNAs. This early system relied on AARS:amino acid recognition and the AARS:proto-tRNA interaction to form the so called operational code.^9^ To improve our understanding of the functional state of this early stage of protein synthesis, we used Class I AARS as a model system. We selected the most conserved parts of the catalytic domain (dubbed the catalytic core) of 10 specialized Class I AARSs to guide structure-based phylogenetic analysis and ancestor reconstruction. The reconstructed early ancestors of the Class I AARS catalytic domain support the hypothesis of statistical primitive translation^11^. Specifically, we found that Anc-All-minus (the ancestor of all class I AARSs except TyrRS and TrpRS, which emerged independently; **Fig. 1**,^33^), as well as Anc-hydrophobic and Anc-charged ancestors (which preceded AARS specialization) have the wide active sites where the determinants that define the amino acid specificity in the contemporary specialized enzymes are replaced by small, uncharged residues (**Fig. 1, Table 1**). Thus, the primitive translation machinery was likely of a generalist type and gradually proceeded towards specialization. Indeed, our data suggest that the earliest ancestor (Anc-All-minus) was capable to select only for L-α-amino acids setting up the chemistry and stereochemistry of the protein alphabet. The next step comprised selection against the amino acid substrate side chains according to the physiocochemical features, starting from the generalist ancestors Anc-hydrophobic and Anc-charged. It has been suggested that some abiotic amino acids – such as norvaline, alpha-aminobutyrate, and ornithine – could have participated in primitive translation ^41–43^ regardless their absence in modern proteins. The apparent broad selectivity of reconstructed early ancestral states corroborates such hypotheses.

Class I AARS phylogeny (**Fig. 1**) shows the enzymes specialized for the late amino acids appearing as early branches at the tree. This unusual feature either contrasts the consensus order of amino acid appearance^34^ or shows some novel feature of enzyme evolution. We opted for the latter and proposed the new model of enzyme emergence, the generalist maintaining model (**Fig. 4**). According to this model, which combines features of both neo-functionalization and sub-functionalization models, the generalist AARS ancestor is kept roughly unchanged after duplications and repeatedly gives rise to specialized AARSs. This model is applicable during the early stages of translation where co-evolution with amino acid takes place and translational fidelity is not a must. Furthermore, it is particularly relevant to complex systems^39^ such as translation because specialization of AARSs could not have occurred independently. Rather, such a system demands co-specialization of cognate tRNAs (*i*.*e*., addition of the anticodon stem), as well as coevolution of the ribosome that decoded these tRNAs and of the mRNAs that encoded proteins at any given stage. Thus, the ancestral generalist system could not be abandoned before an entire specialized translation system evolved, including the acquisition of tRNA binding elements of AARSs (the anticodon domains).

## METHODS

### Structure-based sequence alignments

The available non-redundant crystal structures of Class I AARSs enzymes were examined by scanning the relevant ECOD HUP domain F-groups (2005.1.1.82/139/140/141/143/144/147/150/220). Representative structures were selected for further analysis using the following criteria: (*i*) prokaryote enzymes, when available both a bacterial and archaeal representative, and a eukaryote enzyme when no structure of an archaeal enzyme is available); (*ii*) structures with a bound analog of the aminoacyl-AMP intermediate (or only the amino acid substrate if a complex with the intermediate is unavailable). GlnRS that was shown to have diverged from GluRS and has an essentially identical core structure was not included. Overall, 24 structures were selected and their various domains, including the HUP domain that corresponds to the synthetic domain, and the CP and anticodon binding domains, were catalogued using the original ECOD assignments with some manual refinements (**Table S1**). Seven structures of HIGH Clade 2 enzymes that represent the key families of this clade^28^ were used as outgroup (listed in **Table S2**). The HUP domains of these 31 structures were examined and segments that are not shared by all structures, or are present yet could not be aligned, were removed. The resulting core domains were aligned using PyMol (the PyMol pse file is provided as **Dataset S1**) to the core of the MetRS (1pg0) structure. The pairwise RMSD values for the aligned cores (Table S3) support selection of MetRS (1pg0) as a prototypic core. Overall, the alignable core was divided to five segments (β1α1β2, α2, α3, β4α4, β5) that included all the major functional elements except the so-called KMSKS loop (**Fig. 2, Fig. S3**). Although the sequence of this loop that extends from β5 is relatively conserved, its conformation is highly diverse and hence this loop could not be aligned. The segment that corresponds to this loop was later added to the seed sequence alignment (see below). A sequence alignment was then generated of the core segments of the 31 aligned structures. This ‘seed’ alignment was manually refined ensuring that aligned residues occupy the same structural position (rather than maximizing sequence overlaps as in automated sequence alignments). Both the backbone location and the side chain orientation were considered. Specifically, we ensured that residues whose side chains point into the core and/or the active site occupy the same position in the alignment, and conversely that those whose side chains point to the domain’s exterior align. The KMSKS loop was added to the last segment (segment 7) and this segment was aligned based on sequence only (the ‘seed’ alignment is shown in Fig. S3, and provided as fasta file as part of Dataset S2).

Next, using the Pfam HMM profiles, sequences of the HUP domains that belong to the Class I AARSs families were retrieved from a set of 738 representative archaeal and bacterial genomes that span the prokaryote tree of life^30^ using HMMsearch with the Pfam-defined gathering threshold^44^. These sequences were further clustered at 50% identity with CD-HIT to remove redundancy^45^. Overall, 531 sequences were taken, and these were aligned to the ‘seed’ alignment while ‘freezing’ the latter using Mafft with the options --*add* and -- *keeplength* (the alignment’s fasta file is provided in **Dataset S2**). In parallel, an automated alignment of the entire HUP domain (with no division to segments) was generated using Mafft with the option --*linsi* (provided in fasta format as **Dataset S4**) and automatically trimmed with trimAl to remove gappy regions. In this full-length automated alignment, most positions aligned also in a structurally relevant manner, and especially those in the conserved segments (foremost in β1-α1-β2, but also in many positions in the other segments). However, several residues that were well aligned sequence-wise were found to occupy different positions in the structure. Accordingly, our analysis relied foremost on the core segments alignment.

### Phylogenetic trees and ancestral inference

Phylogenetic trees were inferred from both the core segments alignment and the catalytic domain full-length automated alignment, using FastTree, RaxML and PhyML all with Jones-Taylor-Thornton (JTT) evolutionary models^46–48^ (provided in Newick format as **Dataset S3** and **Dataset S5**). The trees were rooted with a HUP outgroup. The HUP clade placement differs across trees being either an outgroup (PhyML and FastTree) or clustering together with TrpRS (RaxML). Although the trees differ as shown in **Fig. S4**, they converged regarding the key nodes. Ancestral inference was performed on the tree generated by RaxML derived from the core segments alignment that is likely to be more reliable. Ancestral states were inferred using codeml from the PAML package^49^. Ancestral sequences and a phylogenetic tree annotated with ancestral nodes are provided in **Datasets S6, S7**, respectively.

## Acknowledgment

Dan S. Tawfik passed away at the time of manuscript finalization. This work was supported by the Swiss Enlargement Contribution in the framework of the Croatian-Swiss Research Programme, Grant IZHRZ0_180567 to I.G.-S., and by Grant 94747 of the Volkswagen Foundation to D.S.T. D.S.T. was the Nella and Leon Benoziyo Professor of Biochemistry.

## Supplementary information

Supplementary Information (SI Appendix) is available online.

