## Supplementary File 1 for "The evolutionary history of class I aminoacyl-tRNA synthetases indicates early statistical translation"

Department of Chemistry,

Faculty of Science,

University of Zagreb,

Zagreb, Croatia

Liam M. Longo

Earth-Life Science Institute

Tokyo Institute of Technology

Tokyo, Japan

**Supplementary data consist of:**

Supplementary Materials and Methods

Supplementary Figures (1 to 5)

Supplementary Tables (1 to 3)

Datasets:

Dataset S1: Structural alignment of Class I core segments (PyMOL session)

Dataset S2: Structure-guided sequence alignment of Class I core segments (FASTA file)

Dataset S3. Phylogenetic trees calculated from the structure-guided sequence alignment of Class I core segments (Newick files)

Dataset S4: Automated alignment of Class I synthetic domains (FASTA file)

Dataset S5: Phylogenetic trees calculated from the automated alignment of the Class I synthetic domains (Newick files)

Dataset S6: Phylogenetic trees with annotated ancestral nodes (Newick files)

Dataset S7: Alignment of the ancestral sequences and the core segments (FASTA file)

**Supplementary Materials and Methods**

Evolutionary Analysis of Class II AARSs

The analysis pipeline used for Class II AARSs closely followed that of Class I AARSs, as described in the main text. Briefly, 28 representative Class I AARS crystal structures from the relevant ECOD F-groups (314.1.1.5/8/9/12/13/14/21/57) were selected (**Fig. S1C and S1D**). Priority was given to i) prokaryotic enzymes, including both bacterial and archaeal representatives and ii) structures bound to small ligands, preferably an aminoacyl-adenylate intermediate or its sulfamoyl analogue. Of the 28 representative structures chosen, only four were unliganded. If the structure of an archaeal representative was missing, a eukaryotic one was selected instead. Using these representatives, conserved regions of secondary structure – that is, the Class II AARS core – were identified (α0, β1, α1, β2, β3, β4, β5, β6, β7, see **Fig. S1**) and found to include the characteristic Class II AARS motifs 1 to 3 ^1^. The core structures were then superimposed onto the core of prolyl-tRNA synthetase (ProRS; PDB code: 2j3l) using PyMOL (Schrödinger & DeLano). The resulting structure alignment was used to guide a sequence alignment of the core domains (referred to here as the ‘seed’ alignment).

Using HMMsearch with HMM profiles from Pfam (PF13393, PF01409, PF00587, PF01411, PF00152, PF02091) and the Pfam-defined gathering threshold, 6682 sequences of Class II AARSs were retrieved from a set of 738 representative bacterial and archaeal proteomes^2^. CD-HIT^3^ was then used to apply a 75% identity cutoff. The newly gathered sequences were then added to the structure-guided seed alignment by MAFFT using the options *–add* and *–keeplength.*^4^ Phylogenetic trees of the resulting alignment were calculated using FastTree^5^, RaxML^6^ and PhyML^7^. In contrast to Class I AARS, which has the well-defined outgroup ATP/CTP-dependent ligases (HIGH-Clade 2)^8^, the outgroup of Class II AARS is elusive. Although biotin protein ligase (ECOD F-group BPL_LplA_LipB; PDB code: 1x01; **Fig. S1B**) is abundant and structurally similar^9,10^ to Class II AARSs, the mode of ligand binding and the sequences are too diverged to generate a reliable alignment. Therefore, phylogenetic trees were rooted at the midpoint or by minimal ancestral deviation (MAD) (**Fig. S2**).

**Supplementary Figures**

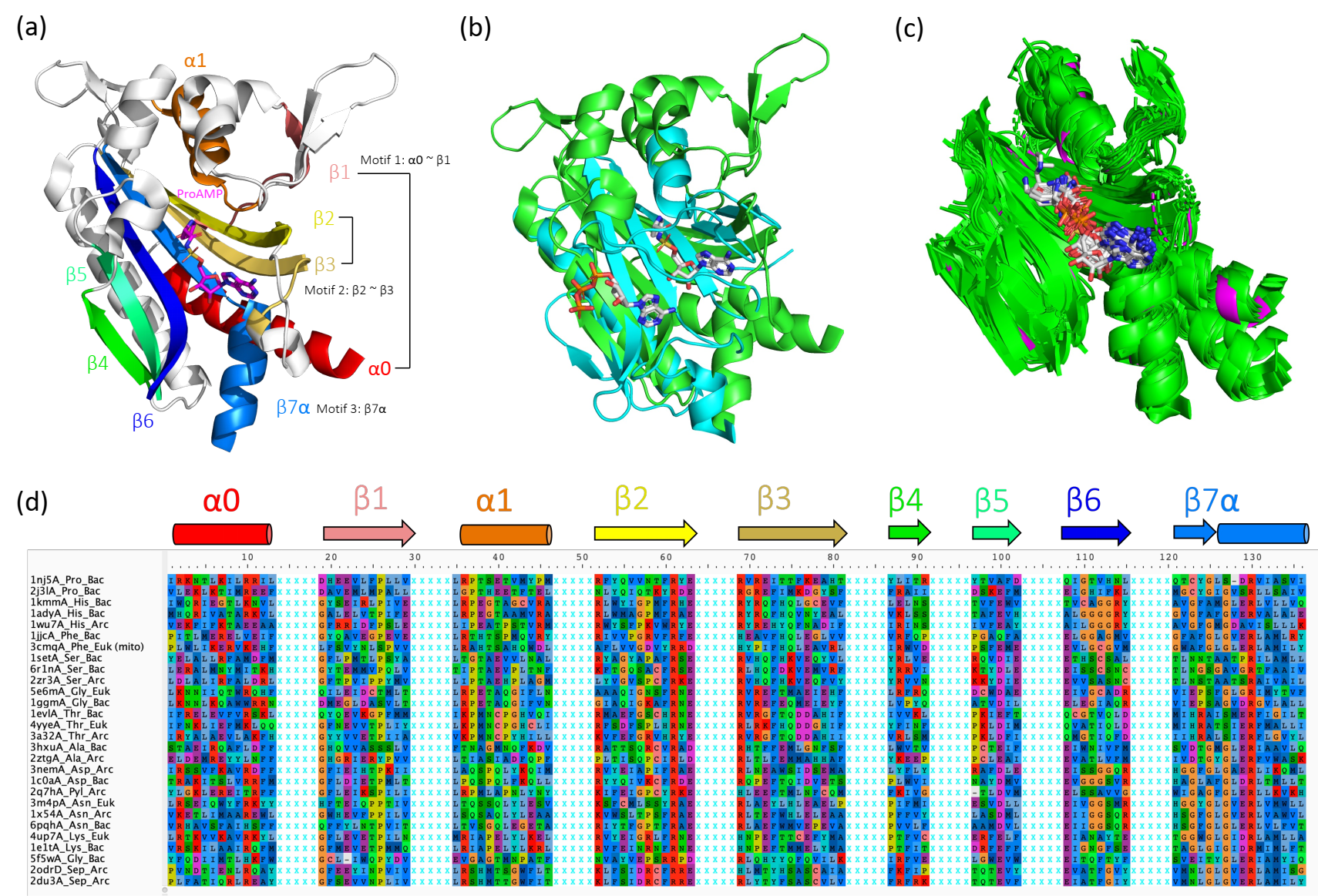

**Figure S1**. (a) Structural highlights of the Class II AARS catalytic domain (illustrated by ProRS, PDB code: 2j3l). The canonical Class II motifs – 1, 2, and 3 – are involved in dimerization, substrate binding and catalysis and are retained in the Class II AARS core. (b) Class II AARSs adopt a unique fold^9,10^ that is shared only with biotin protein ligase. Structural alignment of biotin protein ligase bound to ATP (cyan, PDB code: 1x01) and ProRS bound to an analogue of prolyl-AMP (green), however, shows that the structural features and binding modes of the two proteins have diverged significantly. Consequently, it was not possible to use biotin protein ligase as an outgroup for a structure-guided sequence alignment. Unfortunately, alternative rooting approaches (midpoint and MAD) did not result in a stable reconstruction of the Class II phylogeny (**Fig. S2**) (c) Structural superposition of the Class II AARS catalytic cores against ProRS (magenta) reveals a high degree of structural conservation. The greatest deviation was seen for α0 helix. (d) The structure-guided seed alignment of the Class II catalytic cores. Discontinuous segments are separated by five Xs for clarity. The associated secondary structure elements for each fragment, annotated above the sequence alignment, are colored as in Panel a). Sequences are named as follows: (PDB code)_(AARS type)_(domain of life).

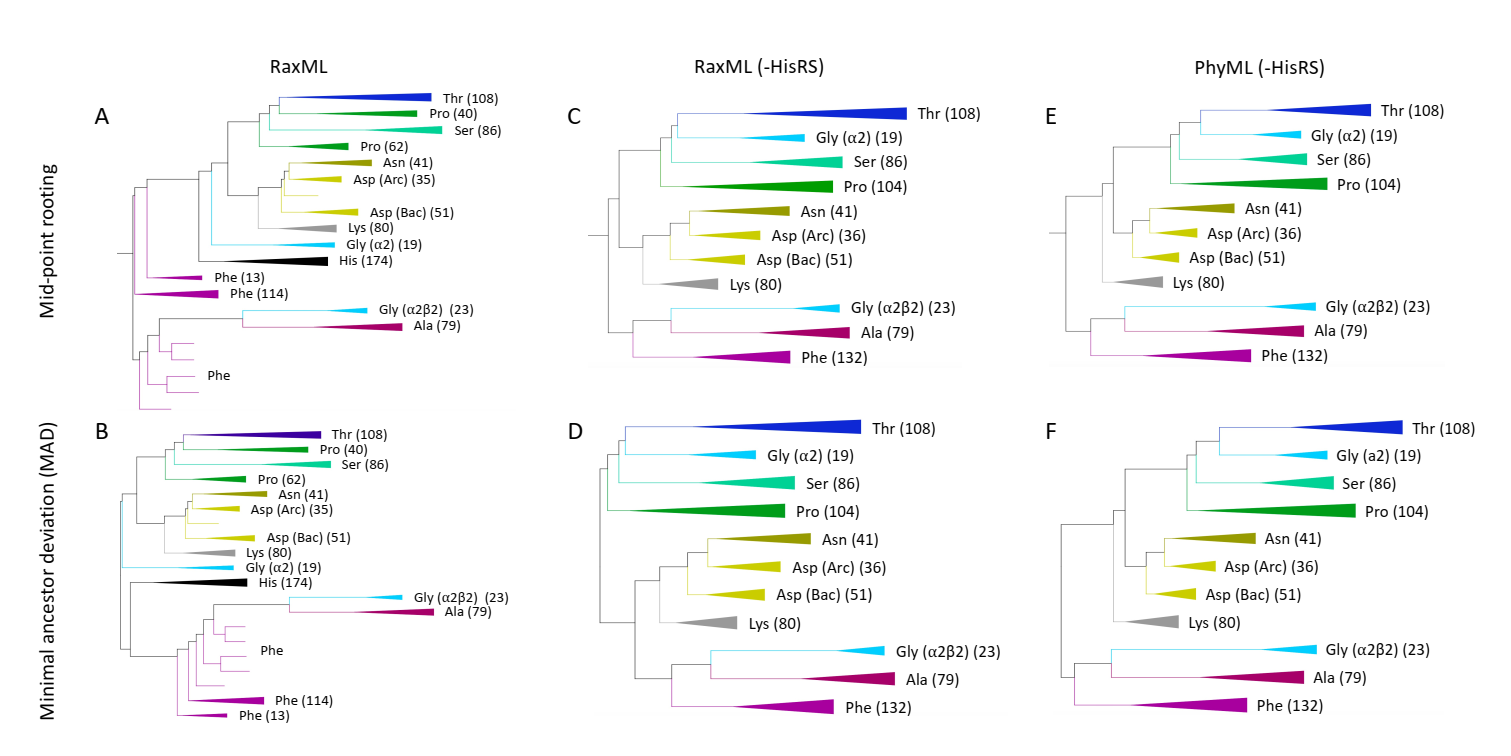

**Figure S2**. Phylogenetic trees of the representative, non-redundant sequences (see **Supplementary Materials and Methods** for the sequence collection process) from ten major Class II AARS families using different rooting methods (midpoint rooting for **A**, **C**, **E** and minimal ancestor deviation rooting for **B**, **D**, **F**) and tree building methods (**A-D** by RaxML, **E-F** by PhyML). Numbers in parentheses are the number of representative sequences within the collapsed clades. Note that GlyRS is polyphyletic^11^: The heterotetrameric α2β2 GlyRS is found in some bacteria, archaea, and eukaryotes, whereas the homodimeric α2 GlyRS is distributed mostly in bacteria. HisRS is excluded in **C-F** to demonstrate that Class II AARS phylogenetic trees are unstable when HisRS is included in the analysis, suggesting that the origin of HisRS is yet to be confidently assigned. The consistency of the three main subtrees and their internal structure – namely, (1) ThrRS, GlyRS (α2), SerRS, ProRS, (2) AsnRS, AspRS, LysRS, and (3) AlaRS, GlyRS (α2β2), PheRS – agrees with the traditional classification of Class II AARSs, denoted IIA, IIB, and IIC, respectively^10^. However, the inconsistency of the phylogeny between the three subtypes when using different phylogenetic analysis methods (**C-F**) precludes a robust ancestral sequence reconstruction of Class II AARSs.

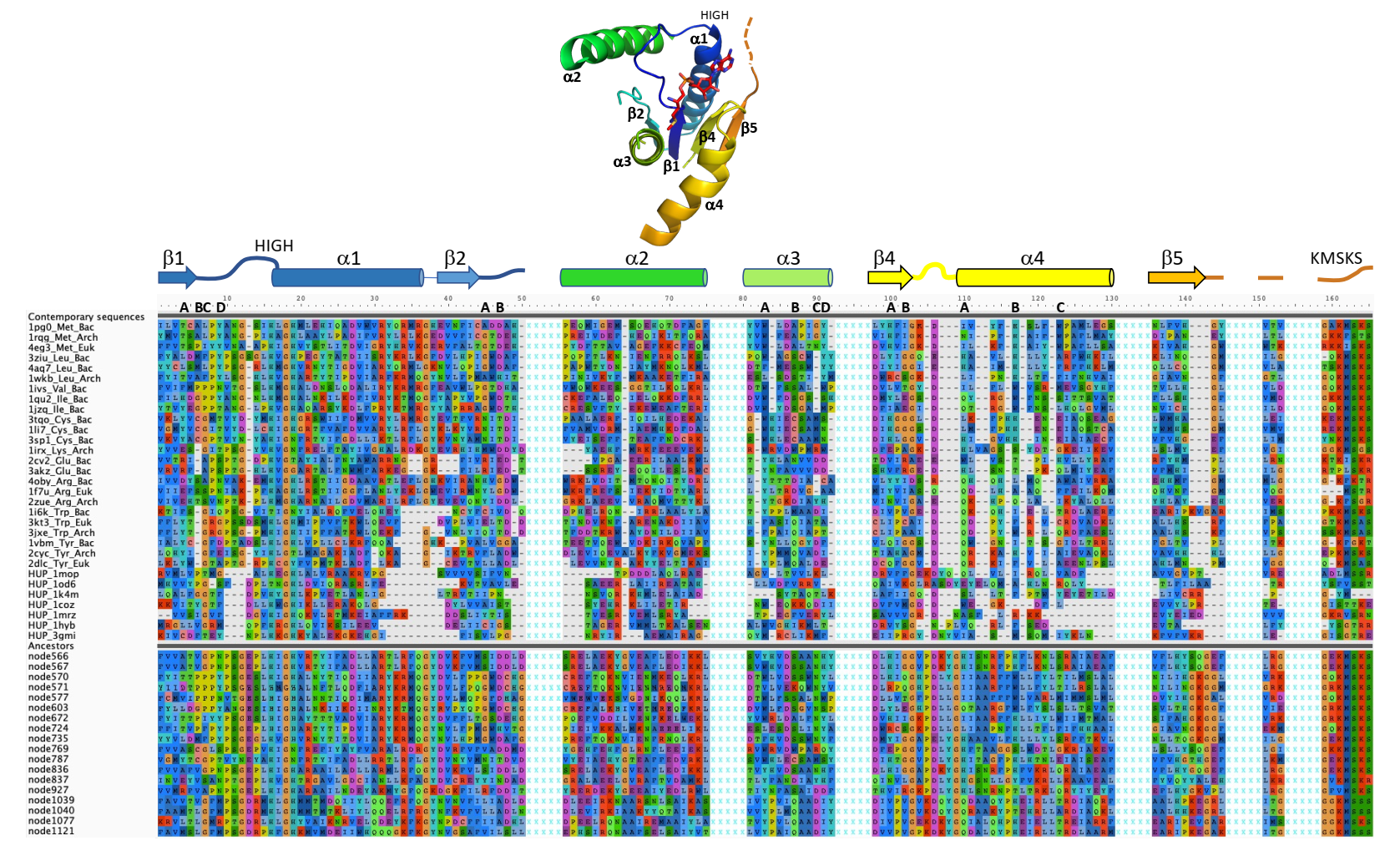

**Figure S3**: A structure-guided seed alignment of the core segments to which the selected reconstructed ancestral nodes were added. The alignment comprises 24 Class I AARS structures (listed in **Table S1**), 7 HIGH-Clade-2 structures (listed in **Table S2**) and 18 reconstructed ancestral nodes using PAML (listed in **Table 1**). The core segments are denoted above the sequence alignment and also shown in a cartoon representation of MetRS (PDB code: 1pg0) that was used as template for assigning these segments. Denoted also are the residues that comprise the amino acid substrate binding pocket (in the residue number ruler). The color and letter codes (A, B, etc) relate to specific secondary structural elements (e.g., β1A, β1B) and follow **Table 1** in the main text. The structural alignment is provided as **Dataset S1**, and the FASTA versions of the sequence alignments shown here are provided in **Datasets S2** and S**7**.

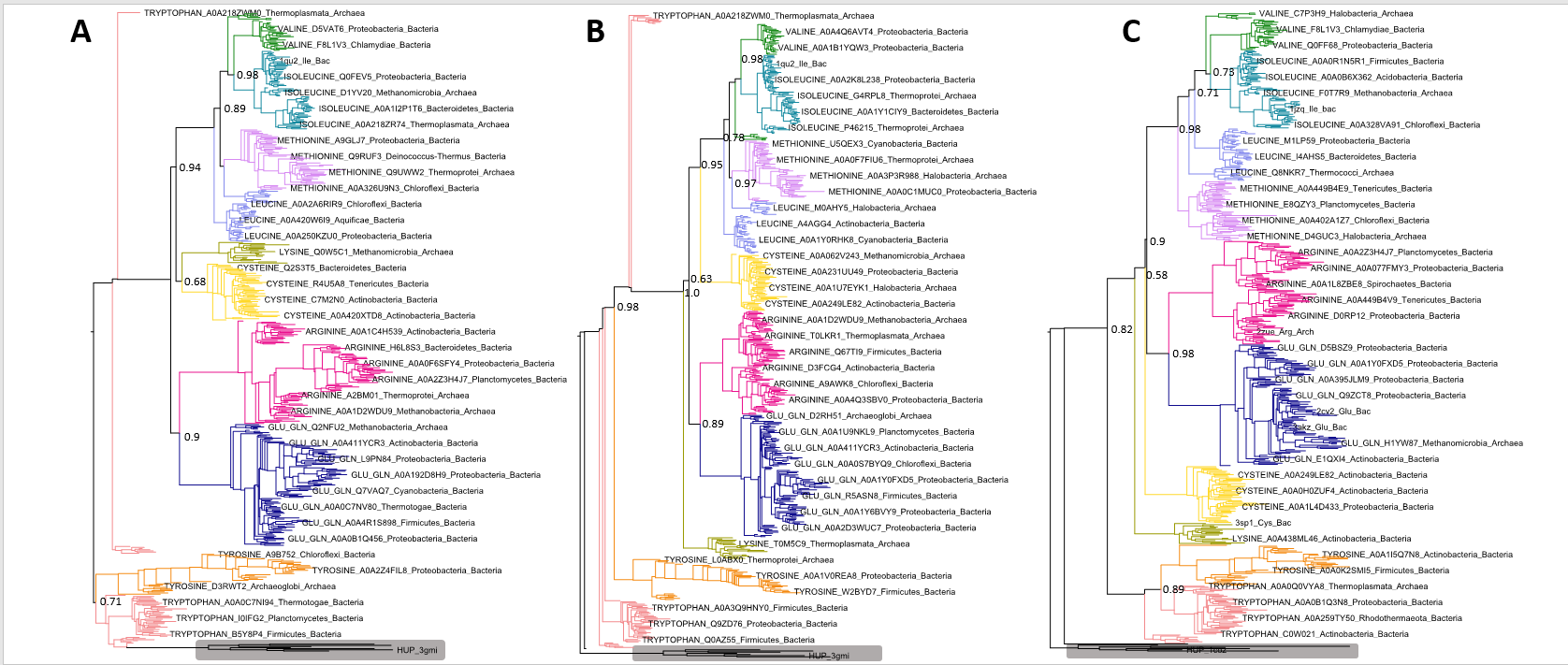

**Figure S4**. The phylogenetic trees of Class I AARS based on a sequence alignment of the AARS core segments generated with three alternative tree-building methods. A) Raxml, B) Phyml, C) FastTree. In all trees, the key confidence values are displayed. Only the splits between CysRS and LysRS may have lower bootstrap values. The HUP outgroup is shaded grey. Raw Newick files of these trees can be found in **Dataset S3**.

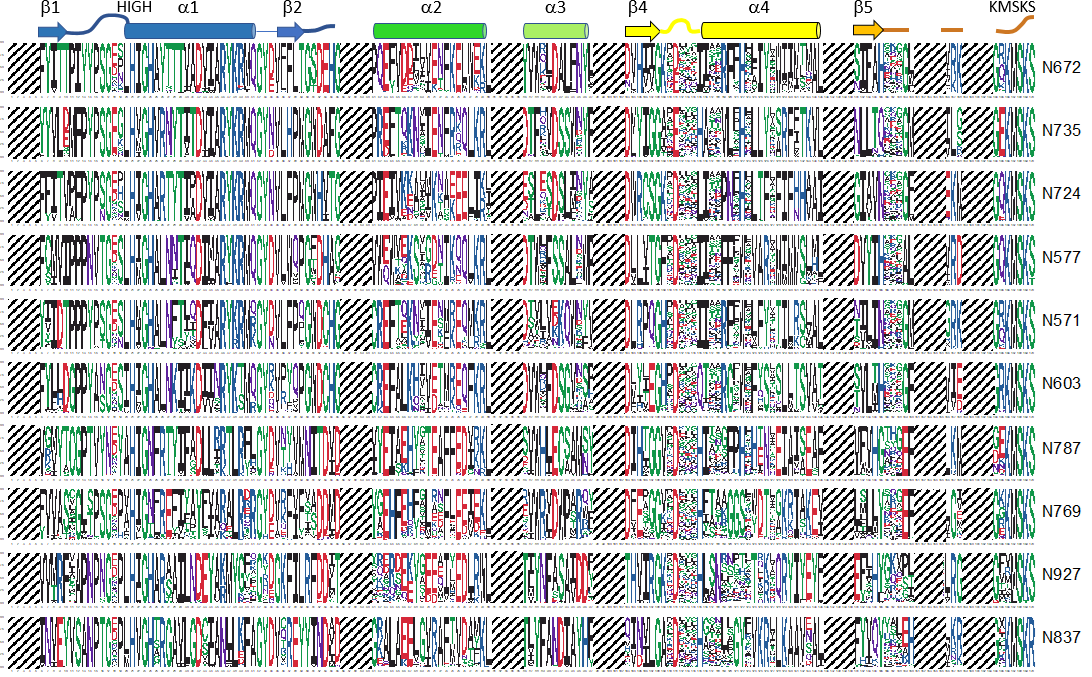

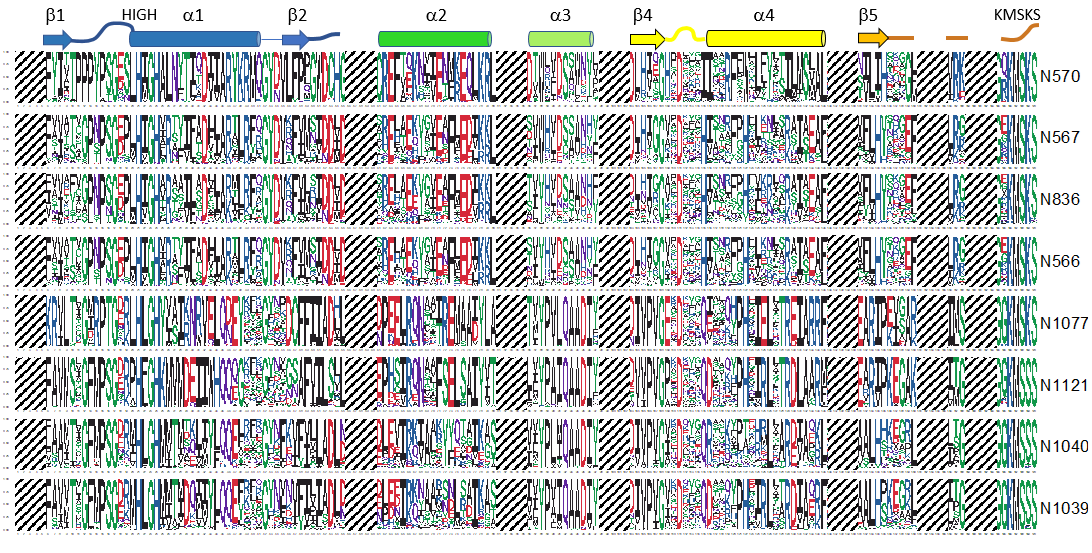

**Figure S5**. Sequence logos of the ancestral states listed in **Table 1**. The most probable ancestral sequences obtained by PAML are given in **Fig. S3** and **Dataset S7**.

**Table S1. AARS structures used for structural alignment**. AARS primary sequences are divided according to the enzyme structural elements and domains. For domains classified by ECOD, a code consisting of four numbers separated by periods is given, representing the X, H, T, and F levels of evolutionary relatedness, respectively. Domains belonging to the same H-group share a common origin and are shaded by the same color. Annotations are done in accordance with ECOD^12^, or manually when required.

| **PDB ID**  **species**  **family** | **N-terminal extension** | **HUP^[1]^**  **1^st^ part** | **H_B3_^[2]^** | **CP1**  **1^st^ part^[3]^** | **Editing domain** | **CP1**  **2^nd^ part** | **CP2^[4]^** | **HUP**  **2^nd^ part** | **SC-fold containing**  **KMSKS motif^[5]^** | **C-terminal anticodon binding domain** | **C-terminal**  **(sub)domain 1** | **C-terminal (sub)domain 2** |
| --- | --- | --- | --- | --- | --- | --- | --- | --- | --- | --- | --- | --- |
| 1 pg0  *E. coli*  MetRS | 1-4^[6]^ | 4-97  2005.1.1.144^[7]^ | 99-116  (17)^[8]^ | 117-230 (113) | none | none | 231-252  (21) | 253-325 | 325-372 | 373-550  140.1.1.1 |  |  |
| 4eg3  *T. brucei*  MetRS^[9]^ | not solved | 239-333  2005.1.1.144 | 334-352  (18) | 354-446  (92) | none | none | 447-470  (23) | 471-547 | 548-593 | 603-767  140.1.1.26 |  |  |
| 1rqg  *P. abyssi*  MetRS | 1-3 | 4-96  2005.1.1.144 | 97-114  (17) | 115-229  (114)  375.1.1.461 | none | none | 230-250  (20) | 251-335 | 336-383 | 384-606  140.1.1.11 |  |  |
| 1qu2  *S. auerus*  IleRS | 1-50 | 51-155  2005.1.1.147 | 156-174  (18) | 175-200  (25)  375.1.1.84 | 201-39  1.1.1.28 | 395-452  (57)  375.1.1.84 | 453-525  (72)  375.1.1.64 | 526-586 | 586-634 | 634-790  140.1.1.14 | 790-880  3008.1.1.2 | 881-916  377.1.3.1  *anticodon binding* |
| 1jzq  *T. thermophilus* IleRS | 1-39 | 40-150  2005.1.1.147 | 151-170  (19) | 171-196  (25)  375.1.1.84 | 197-387  1.1.1.28 | 388-445  (57)  375.1.1.84 | 446-515  (69)  375.1.1.65 | 516-583 | 584-630 | 631-794  140.1.1.14 | not solved | not solved |
| 1ivs  *T. thermophilus* ValRS | 1-34 | 35-145  2005.1.1.147 | 146-162  (16) | 166-190  (24)  375.1.1.84 | 193-339  1.1.1.31 | 342-401  (59)  375.1.1.84 | 402-452  (50)  375.1.1.48 | 454-520 | 521-567 | 568-711  140.1.1.14 | 712-796  3008.1.1.1 | 797-862  192.7.1.6  *anticodon binding* |
| 4aq7  *E. coli*  LeuRS | 1-33 | 34-127  2005.1.1.147 | 128-147  (19) | 148-225  (77)  375.1.1.84 | 226-415  1.1.1.28 | none^[10]^ | 416-490  (74)  375.1.1.387 | 491-568 | 569-661^[11]^  375.14.1.1 | 662-796  140.1.1.14 | 797-860  221.15.1.1  *binds variable*  *arm and D-loop* |  |
| 3ziu  *M. mobile* LeuRS | 1-36 | 37-132  2005.1.1.147 | 133-150  (17) | 151-249  (98)  375.1.1.84 | none^[12]^ | none | 250-318  (78)  375.1.1.452 | 319-402 | 403-449 | 450-575  140.1.1.14 | 577-637  221.15.1.1 |  |
| 1wkb  *P. horikoshii*  LeuRS | 1-34 | 35-155  2005.1.1.147 | 156-173  (17) | 176-205  (29)  375.1.1.257 | 206-443  1.1.1.31 | 444-507  (63)  375.1.1.257 | 508-526  (18)  375.1.1.262 | 527-643 | 644-689 | 690-810  140.1.1.11 | 810-967  not solved |  |
| 1li7  *E. coli*  CysRS | 1-21 | 22-112  2005.1.1.144 | 113-131  (18) | 133-181^[13]^  (48)  375.1.1.466 | none | none | 182-203  (21)  375.1.1.466 | 204-258 | 259-304 | 305-402  140.1.1.16 | 403-461^[14]^  not solved  *anticodon binding* |  |
| 3sp1  *B. burdorferi* CysRS | 1-21 | 22-125  2005.1.1.141 | 126-141  (15) | 142-189  (47)  375.1.1.387 | none | none | 190-215  (25) | 216-271 | 272-316 | 317-426^[15]^ | 431-480 |  |
| 3tqo  *C. burnetti*  CysRS | 1-22 | 23-113  2005.1.1.144 | 114-132  (18) | 133-181^[16]^  (48)  375.1.1.459 | none | none | 182-204  (22)  375.1.1.459 | 205-260 | 261-305 | 306-403  140.1.1.16 | 404-460  not solved |  |
| 1irx  *P. horikoshii* LysRS | 1-20 | 21-125  2005.1.1.143 | 126-162^[17]^  (36) | 170-206^[18]^  (36)  375.1.1.374 | none | none | 207-215  (8)  375.1.1.374 | 217-270 | 271-318 | 318-422  101.8.1.8 | 441-523 |  |
| 1f7u  Yeast  ArgRS | 2-135  310.1.1.1  tRNA (D loop) binding domain | 136-294  2005.1.1.141 | 295-311  (16) | 315-334  (19)^[19]^ | none | none | 335-345  (10) | 346-404 | 405-473^[20]^ | 474-607  140.1.1.15 |  |  |
| 2zue  *P. horikoshii*  ArgRS | 1-119  310.1.1.1 | 120-269  2005.1.1.141 | 270-288  (18) | 289-318  (29) | none | none | 319-329  (10) | 330-417 | 418-492 | 493-629 |  |  |
| 4oby  *E. coli*  ArgRS | 1-107  310.1.1.1 | 108-256  2005.1.1.141 | 257-274  (17) | 275-299  (24) | none | none | 300-311  (11) | 312-371 | 372-444 | 445-576  140.1.1.15 |  |  |
| 3akz  *T. maritima*  GluRS | 1-25 | 26-103  2005.1.1.140 | 104-121  (17) | 122-185^[21]^  (63)  8002.1.1.1 | none | none | 186-195  (9)  8002.1.1.1 | 196-246 | 247-315  8002.1.1.1 | 316-386^[22]^  101.8.1.3 | 387-486  *anticodon binding* |  |
| 2cv2  *T. thermophilus* GluRS^[23]^ | none | 1-80  2005.1.1.140 | 81-99  (18) | 100-173  (73)  8002.1.1.1 | none | none | 174-185  (9) | 186-235 | 236-304 | 305-374  101.8.1.1 | 375-468 |  |
| 2dlc  Yeast  TyrRS | 1-39 | 40-123  2005.1.1.139 | 124-153^[24]^,^[25]^  (29) | 154-162^[26]^ | none | none | none | 163-218 | 219-241^[27]^ | 242-394^[28]^ |  |  |
| 1vbm  *E. coli*  TyrRS | 1-39 | 40-125  2005.1.1.139 | 126-163  (37) | 164-167 | none | none | none | 168-227 | 228-251 | 252-322^[29]^ | 322-424^[30]^  not solved  *variable arm binding* |  |
| 2cyc  *P. horioshii*  TyrRS | 1-33 | 34-120    2005.1.1.139 | 121-154  (33) | 155-161 | none | none | none | 162-225 | 226-264 | 265-375^[31]^ |  |  |
| 3kt3  Yeast  TrpRS | 1-102^[32]^ | 103-183  2005.1.1.82 | 184-214  (30) | 215-219 | none | none | none | 220-285 | 286-309 | 310-432^[33]^ |  |  |
| 3jxe  *P. horikoshii* TrpRS | 1-73 | 74-157  2005.1.1.139 | 158-182  (24) | 183-187 | none | none | none | 188-243 | 244-267 | 268-386 |  |  |
| 1i6k  *G.stearothermophilus* TrpRS | none^[34]^ | 1-80  2005.1.1.150 | 81-113  (32) | 114-117 | none | none | none | 118-182 | 183-208 | 209-328 |  |  |

**Table S2. List of HIGH-NPTs-Clade 2 structures used as outgroup**

| PDB | Enzyme name; organism | HUP domain ECOD annotation | ECOD F-group | Reference structure in HUP analysis |
| --- | --- | --- | --- | --- |
| 1od6 | phosphopantetheine adenylyltransferase; *T. thermophilus* | [e1od6A1](http://prodata.swmed.edu/ecod/complete/domain/e1od6A1)  [A:1-160] | CTP_transf_like  2005.1.1.218 |  |
| 1k4m | nicotinate mononucleotide adenylyltransferase; *E. coli* | [e1k4mC1](http://prodata.swmed.edu/ecod/complete/domain/e1k4mC1)  [C:1-213] | CTP_transf_like  2005.1.1.218 |  |
| 1hyb | Nicotinamide mononucleotide adenylyltransferase; M. thermoautotrophicum | [e1hybA1](http://prodata.swmed.edu/ecod/complete/domain/e1hybA1)[A:4-170] | CTP_transf_like  2005.1.1.218 | As 1od6 % 1k4m above |
| 1coz | cytidylyltransferase: CTP:glycerol-3-phosphate cytidylyltransferase; B. subtilis | [e1cozA1](http://prodata.swmed.edu/ecod/complete/domain/e1cozA1)  [A:1-126] | CTP_transf_like_1  2005.1.1.6 |  |
| 3gmi | unknown function; M. jannaschii | [e3gmiA1](http://prodata.swmed.edu/ecod/complete/domain/e3gmiA1)  [A:1-271] | HIGH_NTase1_ass_N 2005.1.1.134 |  |
| 1mop | pantothenate synthetase; *M. tuberculosis* | [e1mopA3](http://prodata.swmed.edu/ecod/complete/domain/e1mopA3)  [A:3-190] | Pantoate_ligase_N 2005.1.1.146 | See 2a86A* |
| 1mrz | flavin-binding protein; T. maritima | [e1mrzA2](http://prodata.swmed.edu/ecod/complete/domain/e1mrzA2)  [A:2-158] | FAD_syn  2005.1.1.133 | See 1t6yB* |

**Table S3.** **Structural Alignment RMSD (Å) for Class I AARS and HUP Cores in Dataset 1.**

|  | 1ivs_Val_bac | 1jzq_Ile_bac | 1qu2_Ile_bac | 1pg0_Met_bac | 1rqg_Met_arch | 4eg3_Met_euk | 4aq7_Leu_bac | 3ziu_Leu_bac | 1wkb_Leu_arch | 1irx_Lys_arch | 3sp1_Cys_bac | 3tqo_Cys_bac | 1li7_Cys_bac | 4oby_Arg_bac | 2zue_Arg_arch | 1f7u_Arg_euk | 2cv2_Glu_bac | 3akz_Glu_bac | 1vbm_Tyr_bac | 2cyc_Tyr_arch | 2dlc_Tyr_euk | 1i6k_Trp_bac | 3jxe_Trp_arch | 3kt3_Trp_euk | 1mop_hup | 1od6_hup | 3gmi_hup | 1mrz_hup | 1k4m_hup |
| --- | --- | --- | --- | --- | --- | --- | --- | --- | --- | --- | --- | --- | --- | --- | --- | --- | --- | --- | --- | --- | --- | --- | --- | --- | --- | --- | --- | --- | --- |
| 1ivs_Val_bac | 0.0 | 1.6 | 1.2 | 1.5 | 1.6 | 1.5 | 1.2 | 1.3 | 1.1 | 2.1 | 1.8 | 1.7 | 1.7 | 2.4 | 2.6 | 2.5 | 2.0 | 1.9 | 1.9 | 2.1 | 1.9 | 2.0 | 1.8 | 1.7 | 2.8 | 2.7 | 2.3 | 2.9 | 3.1 |
| 1jzq_Ile_bac | 1.6 | 0.0 | 1.3 | 1.9 | 1.9 | 1.8 | 1.6 | 1.7 | 1.6 | 1.9 | 1.9 | 1.9 | 1.9 | 2.4 | 2.7 | 2.5 | 2.0 | 2.0 | 2.4 | 2.3 | 2.2 | 2.2 | 2.1 | 2.0 | 2.5 | 3.0 | 2.6 | 3.2 | 3.4 |
| 1qu2_Ile_bac | 1.2 | 1.3 | 0.0 | 1.5 | 1.6 | 1.6 | 1.4 | 1.4 | 1.5 | 2.3 | 1.6 | 1.6 | 1.5 | 2.4 | 2.7 | 2.4 | 2.0 | 1.9 | 2.1 | 2.1 | 1.8 | 2.2 | 2.1 | 1.9 | 2.8 | 2.9 | 2.3 | 2.9 | 3.0 |
| 1pg0_Met_bac | 1.5 | 1.9 | 1.5 | 0.0 | 0.8 | 0.9 | 1.4 | 1.5 | 1.4 | 2.0 | 1.6 | 1.7 | 1.6 | 2.2 | 2.4 | 2.2 | 1.9 | 1.8 | 1.9 | 1.9 | 1.7 | 2.0 | 1.8 | 1.5 | 2.2 | 2.2 | 2.1 | 2.9 | 3.0 |
| 1rqg_Met_arch | 1.6 | 1.9 | 1.6 | 0.8 | 0.0 | 1.2 | 1.4 | 1.6 | 1.5 | 2.1 | 1.7 | 1.7 | 1.7 | 2.4 | 2.7 | 2.5 | 2.0 | 1.9 | 2.0 | 2.1 | 1.8 | 1.9 | 1.9 | 1.7 | 2.4 | 2.4 | 2.2 | 2.8 | 3.1 |
| 4eg3_Met_euk | 1.5 | 1.8 | 1.6 | 0.9 | 1.2 | 0.0 | 1.3 | 1.4 | 1.4 | 1.8 | 1.7 | 1.7 | 1.6 | 2.3 | 2.6 | 2.4 | 1.7 | 1.7 | 2.0 | 2.0 | 1.9 | 2.0 | 1.7 | 1.6 | 2.1 | 2.4 | 2.1 | 3.1 | 3.2 |
| 4aq7_Leu_bac | 1.2 | 1.6 | 1.4 | 1.4 | 1.4 | 1.3 | 0.0 | 0.9 | 1.1 | 2.0 | 1.8 | 1.7 | 1.7 | 2.5 | 2.8 | 2.6 | 1.8 | 1.8 | 1.9 | 2.2 | 1.9 | 2.0 | 1.8 | 1.6 | 2.3 | 2.6 | 2.2 | 2.9 | 3.1 |
| 3ziu_Leu_bac | 1.3 | 1.7 | 1.4 | 1.5 | 1.6 | 1.4 | 0.9 | 0.0 | 1.4 | 2.2 | 2.0 | 1.9 | 1.8 | 2.6 | 2.9 | 2.7 | 2.0 | 1.9 | 2.1 | 2.3 | 2.0 | 2.2 | 1.9 | 1.8 | 2.4 | 3.0 | 2.2 | 3.0 | 3.1 |
| 1wkb_Leu_arch | 1.1 | 1.6 | 1.5 | 1.4 | 1.5 | 1.4 | 1.1 | 1.4 | 0.0 | 1.8 | 1.5 | 1.5 | 1.6 | 2.5 | 2.6 | 2.5 | 1.7 | 1.7 | 1.8 | 2.1 | 1.8 | 1.7 | 1.7 | 1.6 | 2.2 | 2.3 | 2.3 | 2.8 | 3.1 |
| 1irx_Lys_arch | 2.1 | 1.9 | 2.3 | 2.0 | 2.1 | 1.8 | 2.0 | 2.2 | 1.8 | 0.0 | 1.7 | 2.1 | 1.9 | 1.7 | 2.0 | 2.0 | 1.5 | 1.7 | 2.6 | 2.3 | 2.3 | 2.0 | 2.2 | 2.2 | 1.9 | 2.5 | 2.1 | 2.6 | 3.2 |
| 3sp1_Cys_bac | 1.8 | 1.9 | 1.6 | 1.6 | 1.7 | 1.7 | 1.8 | 2.0 | 1.5 | 1.7 | 0.0 | 0.8 | 0.8 | 2.1 | 2.4 | 2.2 | 1.8 | 1.8 | 2.3 | 2.1 | 1.8 | 2.0 | 2.1 | 1.8 | 2.6 | 2.3 | 2.3 | 3.1 | 3.2 |
| 3tqo_Cys_bac | 1.7 | 1.9 | 1.6 | 1.7 | 1.7 | 1.7 | 1.7 | 1.9 | 1.5 | 2.1 | 0.8 | 0.0 | 0.6 | 2.2 | 2.7 | 2.3 | 1.8 | 1.8 | 2.4 | 2.2 | 1.8 | 2.1 | 2.1 | 1.9 | 2.7 | 2.5 | 2.4 | 3.3 | 3.1 |
| 1li7_Cys_bac | 1.7 | 1.9 | 1.5 | 1.6 | 1.7 | 1.6 | 1.7 | 1.8 | 1.6 | 1.9 | 0.8 | 0.6 | 0.0 | 2.1 | 2.6 | 2.3 | 1.9 | 1.8 | 2.4 | 2.2 | 1.8 | 2.0 | 2.1 | 1.9 | 2.8 | 2.8 | 2.3 | 3.1 | 2.8 |
| 4oby_Arg_bac | 2.4 | 2.4 | 2.4 | 2.2 | 2.4 | 2.3 | 2.5 | 2.6 | 2.5 | 1.7 | 2.1 | 2.2 | 2.1 | 0.0 | 1.6 | 1.2 | 2.0 | 2.1 | 2.8 | 2.4 | 2.4 | 2.4 | 2.4 | 2.4 | 2.1 | 3.0 | 2.2 | 2.9 | 3.1 |
| 2zue_Arg_arch | 2.6 | 2.7 | 2.7 | 2.4 | 2.7 | 2.6 | 2.8 | 2.9 | 2.6 | 2.0 | 2.4 | 2.7 | 2.6 | 1.6 | 0.0 | 1.4 | 2.4 | 2.4 | 2.9 | 2.2 | 2.5 | 2.6 | 2.5 | 2.3 | 2.3 | 2.6 | 2.4 | 2.5 | 3.3 |
| 1f7u_Arg_euk | 2.5 | 2.5 | 2.4 | 2.2 | 2.5 | 2.4 | 2.6 | 2.7 | 2.5 | 2.0 | 2.2 | 2.3 | 2.3 | 1.2 | 1.4 | 0.0 | 2.3 | 2.2 | 2.6 | 2.2 | 2.3 | 2.5 | 2.5 | 2.4 | 2.3 | 2.5 | 2.3 | 2.7 | 3.3 |
| 2cv2_Glu_bac | 2.0 | 2.0 | 2.0 | 1.9 | 2.0 | 1.7 | 1.8 | 2.0 | 1.7 | 1.5 | 1.8 | 1.8 | 1.9 | 2.0 | 2.4 | 2.3 | 0.0 | 0.9 | 2.3 | 2.2 | 2.0 | 1.9 | 2.0 | 1.9 | 2.2 | 2.2 | 2.3 | 2.8 | 2.9 |
| 3akz_Glu_bac | 1.9 | 2.0 | 1.9 | 1.8 | 1.9 | 1.7 | 1.8 | 1.9 | 1.7 | 1.7 | 1.8 | 1.8 | 1.8 | 2.1 | 2.4 | 2.2 | 0.9 | 0.0 | 2.3 | 2.2 | 1.9 | 2.0 | 2.0 | 1.9 | 2.2 | 2.6 | 2.2 | 2.8 | 2.9 |
| 1vbm_Tyr_bac | 1.9 | 2.4 | 2.1 | 1.9 | 2.0 | 2.0 | 1.9 | 2.1 | 1.8 | 2.6 | 2.3 | 2.4 | 2.4 | 2.8 | 2.9 | 2.6 | 2.3 | 2.3 | 0.0 | 1.7 | 1.7 | 1.9 | 1.8 | 1.6 | 2.4 | 2.3 | 2.3 | 2.6 | 2.9 |
| 2cyc_Tyr_arch | 2.1 | 2.3 | 2.1 | 1.9 | 2.1 | 2.0 | 2.2 | 2.3 | 2.1 | 2.3 | 2.1 | 2.2 | 2.2 | 2.4 | 2.2 | 2.2 | 2.2 | 2.2 | 1.7 | 0.0 | 1.2 | 1.9 | 1.6 | 1.5 | 2.2 | 2.2 | 2.2 | 2.3 | 3.0 |
| 2dlc_Tyr_euk | 1.9 | 2.2 | 1.8 | 1.7 | 1.8 | 1.9 | 1.9 | 2.0 | 1.8 | 2.3 | 1.8 | 1.8 | 1.8 | 2.4 | 2.5 | 2.3 | 2.0 | 1.9 | 1.7 | 1.2 | 0.0 | 1.6 | 1.8 | 1.5 | 2.4 | 2.2 | 2.1 | 2.8 | 3.0 |
| 1i6k_Trp_bac | 2.0 | 2.2 | 2.2 | 2.0 | 1.9 | 2.0 | 2.0 | 2.2 | 1.7 | 2.0 | 2.0 | 2.1 | 2.0 | 2.4 | 2.6 | 2.5 | 1.9 | 2.0 | 1.9 | 1.9 | 1.6 | 0.0 | 1.6 | 1.5 | 2.3 | 2.8 | 2.5 | 2.7 | 3.4 |
| 3jxe_Trp_arch | 1.8 | 2.1 | 2.1 | 1.8 | 1.9 | 1.7 | 1.8 | 1.9 | 1.7 | 2.2 | 2.1 | 2.1 | 2.1 | 2.4 | 2.5 | 2.5 | 2.0 | 2.0 | 1.8 | 1.6 | 1.8 | 1.6 | 0.0 | 0.7 | 2.2 | 2.5 | 2.1 | 2.6 | 3.3 |
| 3kt3_Trp_euk | 1.7 | 2.0 | 1.9 | 1.5 | 1.7 | 1.6 | 1.6 | 1.8 | 1.6 | 2.2 | 1.8 | 1.9 | 1.9 | 2.4 | 2.3 | 2.4 | 1.9 | 1.9 | 1.6 | 1.5 | 1.5 | 1.5 | 0.7 | 0.0 | 2.3 | 2.4 | 2.0 | 2.3 | 3.2 |
| 1mop_hup | 2.8 | 2.5 | 2.8 | 2.2 | 2.4 | 2.1 | 2.3 | 2.4 | 2.2 | 1.9 | 2.6 | 2.7 | 2.8 | 2.1 | 2.3 | 2.3 | 2.2 | 2.2 | 2.4 | 2.2 | 2.4 | 2.3 | 2.2 | 2.3 | 0.0 | 2.7 | 2.3 | 2.6 | 2.9 |
| 1od6_hup | 2.7 | 3.0 | 2.9 | 2.2 | 2.4 | 2.4 | 2.6 | 3.0 | 2.3 | 2.5 | 2.3 | 2.5 | 2.8 | 3.0 | 2.6 | 2.5 | 2.2 | 2.6 | 2.3 | 2.2 | 2.2 | 2.8 | 2.5 | 2.4 | 2.7 | 0.0 | 2.0 | 2.8 | 2.7 |
| 3gmi_hup | 2.3 | 2.6 | 2.3 | 2.1 | 2.2 | 2.1 | 2.2 | 2.2 | 2.3 | 2.1 | 2.3 | 2.4 | 2.3 | 2.2 | 2.4 | 2.3 | 2.3 | 2.2 | 2.3 | 2.2 | 2.1 | 2.5 | 2.1 | 2.0 | 2.3 | 2.0 | 0.0 | 2.5 | 3.0 |
| 1mrz_hup | 2.9 | 3.2 | 2.9 | 2.9 | 2.8 | 3.1 | 2.9 | 3.0 | 2.8 | 2.6 | 3.1 | 3.3 | 3.1 | 2.9 | 2.5 | 2.7 | 2.8 | 2.8 | 2.6 | 2.3 | 2.8 | 2.7 | 2.6 | 2.3 | 2.6 | 2.8 | 2.5 | 0.0 | 3.0 |
| 1k4m_hup | 3.1 | 3.4 | 3.0 | 3.0 | 3.1 | 3.2 | 3.1 | 3.1 | 3.1 | 3.2 | 3.2 | 3.1 | 2.8 | 3.1 | 3.3 | 3.3 | 2.9 | 2.9 | 2.9 | 3.0 | 3.0 | 3.4 | 3.3 | 3.2 | 2.9 | 2.7 | 3.0 | 3.0 | 0.0 |

**Table S4. The amino acid substrate-binding pockets in representative Class I AARSs**. Interacting positions are shaded yellow (side chain interactions), blue (α-amino group recognition), and pink (α-carboxyl group recognition). Double shading indicates interactions with multiple parts of the substrate.

| interacting positions | **1pg0**  **MetRS** | **4aq7**  **LeuRS** | **1ivs**  **ValRS** | **1jzq**  **IleRS** | **1li7**  **CysRS** | **1irx**  **LysRS^[35]^** | **3akz**  **GluRS** | **4oby**  **ArgRS** | **1i6k**  **TrpRS** | **1vbm**  **TyrRS** |
| --- | --- | --- | --- | --- | --- | --- | --- | --- | --- | --- |
| β1 A^[36]^ | _^[37]^ | _ | _ | _ | _ | 25Glu | 28Arg^[38]^ | 118Asp | 5Phe | 37Tyr |
| β1 B^[39]^ | 12Ala | 40Met | 41Pro | 45Gly | 28Cys  Zn shell | deletion^[40]^ | deletion | _ | _ | _ |
| β1 C^[41]^ | 13Leu | 41Leu | 42Pro | 46Pro | 29Gly | 27Gly | 30Ala | 121Ala | _ | _ |
| β1 D^[42]^ | _ | _ | 44Asn | _ | _ | 29Thr | _ | 123Asn | _ | _ |
| β2 A^[43]^ | _ | _ | _ | _ | _ | 64Met | 62Arg | 158His | _ | _ |
| β2 B^[44]^ | 52Asp | 79Asp | 81Asp | 85Asp | 68Thr | 66Asp | 64Glu | _ | _ | _ |
| α3 A | 253Trp | 493Phe | 456Trp | 518Trp | 205Trp | 218Trp | 198Tyr | 313Tyr | 125Tyr | 175Tyr^[45]^ |
| α3 B | 256Ala | 496Ser | 459Ser | 521Ser | 209Cys  Zn shell | 222Trp | _ | 317Asp | 129Met^[46]^ | 179Gln |
| α3 C^[47]^ | _ | 499Tyr | _ | _ | _ | 225Arg | _ | _ | 132Asp | 182Asp |
| α3 D | 260Tyr | _ | _ | _ | _ | 226Trp | _ | _ | _ | _ |
| β4 A^[48]^ | _ | _ | _ | _ | _ | 234Glu | _ | _ | 141Val | _ |
| β4 B^[49]^ | _ | _ | _ | 550Glu | _ | 236Ala | 216Arg | 337Ile | 143Val | _ |
| α4 A^[50]^ | _ | 533His | 491Ile | 554Gln | 230Leu | 240His | 220His | 341Gln | 147 Gln | 201Gln |
| α4 B^[51]^ | 301His | 537His | 495Trp | 558Trp | 234His  Zn shell | 246Ser | _ | _ | _ | _ |
| α4 C | _ | _ | _ | _ | 238Glu  Zn shell | 250Gly | _ | _ | _ | _ |

1. ECOD classification: X: HUP domain-like; H and T: HUP domains.
2. α-Helix following the β3 strand of the HUP core (assigned within the 1^st^ part of HUP in ECOD).
3. The CP1 and CP2 domains are separately classified in ECOD only for IleRS, ValRS, and LeuRS.
4. By ECOD and our classification, the CP1 domain comprises β1 to β3 of the CP core and the CP2 domain extends from β3 to α3. The latter belongs to the 2^nd^ part of the HUP domain.
5. SC fold adopts a β-loop-α-α-β topology. The loop contains the KMSKS motif. The subsequent α-helix, previously assigned to the SC-fold^13^ is assigned here to the helix bundle anticodon binding domain. The SC-fold is not classified by ECOD as a separate domain.
6. Numbers correspond to amino acid positions in the PDB primary sequence.
7. ECOD F-group. If missing, the domain is not (separately) classified in ECOD.
8. Segment length is given in brackets.
9. The N-terminally truncated version was crystallized. Trypanosomatid MetRSs have at their N-terminus a GST-like domain that may participate in protein-protein interactions (Carrot JBC 2021)
10. In bacterial LeuRSs, the editing domain is inserted after the CP1 domain.
11. This segment in *E. coli* LeuRS contains a leucine-specific domain and the KMSKS loop is at the end of the fragment.
12. Naturally lacks the editing domain.
13. The ECOD classification is related to the CP domain containing both CP1 and CP2 parts.
14. Not visualized in this structure. The domain was solved in the presence of tRNA (PDB code: 1u0b) and presents the anticodon binding domain. The domain is not classified in ECOD.
15. Domain is not classified in ECOD. Yet, it structurally resembles the analogous, ECOD-classified domain from Ec IleRS (PDB code: 1li7).
16. The ECOD classification is related to the CP domain containing both CP1 and CP2 parts.
17. Instead of one α-helix, this segment is built from a small α-helix followed by a larger one with a kink.
18. The ECOD classification is related to the CP domain containing both CP1 and CP2 parts.
19. Short CP1 and CP2 built only from the β-strands and loops.
20. Some ArgRS do not have a signature KMSKS motif.
21. The ECOD classification is related to the CP domain containing both CP1 and CP2 parts.
22. The ECOD classification is related to the both C-terminal anticodon binding domains (residues 316-486).
23. Classified using the database of HMM profiles provided by ECOD.
24. In TyrRS and TrpRS, this segment contains a small α-helix followed by a larger one having one or two kinks.
25. This segment is part of a dimer interface in TyrRS and TrpRS
26. CP domain includes one loop connecting HB3 and α3 of the core of the HUP domain. In previous work, the HB3 was assigned as a part of the CP domain.
27. This fragment in TyRS and TrpRS contains just one loop, thus adopting a significantly different topology than the SC domains of IleRS, ValRS, LeuRS, CysRS, ArgRS, LysRS, and GluRS.
28. Novel topology of the anticodon binding domain containing a β-hairpin.^14^ This domain is not classified in ECOD.
29. Topology of the bacterial anticodon domain is slightly different (missing a β-hairpin) from the topology of the eukaryotic and archaeal domains.
30. Present only in bacterial TyrRS where the variable arm is a tRNA identity element. The structure was solved in the presence of tRNA (1h3e) but the domain is not classified as a separate domain in ECOD.
31. Analogous topology to eukaryotic TyrRSs.
32. N-terminal domain containing a β-hairpin that participates in building the Trp/ATP binding site. The domain is not classified as a separate domain in ECOD.
33. Topology of anticodon binding domains in TrpRSs are similar to the topology of the anticodon binding domain of bacterial TyrRS.
34. Bacterial TrpRS do not have an N-terminal domain.^15^
35. The binding pocket cannot be defined as the only available structure is without ligands.
36. The -3 position relative to β1 C.
37. The residue at this position does not establish a contact in the corresponding AARS.
38. The residue is underlined if it is conserved in the ancestor of the corresponding AARS.
39. The -2 position relative to β1 C.
40. The analogous position is missing in this structure.
41. β1 C is the residue at the tip of β1 (or the loop following β1) which binds the α-amino group of the amino acid substrate via backbone carbonyl interactions and forms the binding pocket for the substrate side chain.
42. The +2 position relative to β1 C.
43. The -2 position relative to β2 B.
44. The conserved Asp that binds the α-amino group of the substrate amino acid in all but ArgRS, TyrRS, and TrpRS.
45. Also Asp81 from β2 loop participates in binding the α-amino group of the substrate.
46. Could be Gln that makes a hydrogen bond to the α-amino group of the substrate.
47. The position -1 relative to α3 D.
48. The position -3 relative to the Gly (GXD motif) at the tip of of β4 that anchors the ribose.
49. The position -1 relative to the Gly (GXD motif) at the tip of of β4 that anchors the ribose.
50. +1 relative to the conserved Asp (GXD motif) that positions ribose.
51. +5 position relative to the conserved Asp that positions ribose (for ArgRS, TrpRS and TyrRS is +6 position).
